# Effect of victim relatedness on cannibalistic behaviour of ladybird beetle, *Menochilus sexmaculatus* Fabricius (Coleoptera: Coccinellidae)

**DOI:** 10.1101/2022.09.30.510267

**Authors:** Tripti Yadav, Omkar, Geetanjali Mishra

## Abstract

Cannibalism is taxonomically widespread and has a large impact on the individuals’ fitness and population dynamics. Thus, identifying how the rates of cannibalism are affected by different ecological cues is crucial for predicting species evolution and population dynamics. In current experiment, we investigated how victim relatedness affects the cannibalistic tendencies of different life stages of ladybird, *Menochilus sexmaculatus*, which is highly cannibalistic. We provided larval instars and newly emerged adults of *M. sexmaculatus* with a choice of sibling, half-sibling and non-sibling conspecific eggs as victim of cannibalism. First victim cannibalised and latency to cannibalise were observed along with total number of victims cannibalised after 24 hours. First preference of victim did not differ with life stages of the cannibals though the number of victims cannibalized did increase with advancement in stage. Percentage of total eggs cannibalised also varied significantly with life stage and victim relatedness. First and second instars tend to cannibalise more percentage of siblings and non-sibling eggs while third instars cannibalised more percentage of non-sibling eggs; fourth instars and adults on the other hand cannibalised highest percentage of eggs irrespective of their relatedness. Insignificant effect of victim relatedness was observed on latency to cannibalise eggs, though it varied significantly with the cannibal’s life stage. Shortest latency to cannibalise was recorded for first instars and longest for adults and second instars. In conclusion, kin recognition and its avoidance is stage-specific, with fourth instar and newly emerged adults being less discriminatory as compared to early stages owing to increased evolutionary survival pressure.

## Introduction

In animal taxa, where parents frequently deposit eggs in clusters in a spatially constrained area, the chances of increased levels of competition between the conspecifics is high (Singh *et al*., 2019; Zaviezo *et al*., 2019; Uveges *et al*., 2021). In such a scenario, discrimination between related individuals becomes essential. The presence of kin recognition and kin discrimination during intensive conspecific interactions has been reported in animals ranging from bacteria to vertebrates (Wall, 2016; Henkel and Setchell, 2018; Kalamara *et al*., 2018; Mathiron *et al*., 2019; Anten and Chen, 2021).

Organism responsiveness towards the related individuals can have a major impact on its inclusive fitness (West and Gardner, 2013). Thus, in species where individuals can detect the variations in relatedness (kin and non-kin), behavioural variations can be observed. Relatedness is usually assessed via either phenotypic cues signalling a presence of specific shared genes or genotypes, or contextual cues (Penn and Frommen, 2010; Chung *et al*., 2020;). Sibling cannibalism has been reported across diverse taxa including invertebrates (Chiu *et al*., 2010; Miranda *et al*., 2011), arthropods (Johnson *et al*., 2010; Modanu *et al*., 2014), fishes (Liu *et al*., 2017; Pereira *et al*., 2017), amphibians (Walls and Blaustein, 1995; Park *et al*., 2005; Dugas *et al*., 2016), and birds (Bortolotti *et al*., 1991; Soler *et al*., 2022). In contrast, several studies have reported the identification and avoidance of sibling cannibalism across taxa (Dobler and Kolliker, 2010; Schutt, 2017); these organisms avoid cannibalising related but readily cannibalise unrelated young ones.

A varied range of behavioural and life-history phenotypes appear to have evolved, especially in parts where intense sibling competition occurs (Pfennig and Collins, 1993; Pfennig, 2021). Certain protozoans (Rosati *et al*., 1988; Tollrian and Harvell, 1999), rotifers (Gilbert, 2017), nematodes (Lightfoot *et al*., 2021), insects (Pener and Simpson, 2009), and amphibian larvae (Pfennig and Collins, 1993; Pfennig *et al*., 1993; Pfennig *et al*., 1994) exist as one of two structurally and behaviourally distinct morphs, *i*.*e*. cannibalistic or non-cannibalistic, depending on the environmental conditions they are raised in (Levis and Ragsdale, 2022; Pfennig, 2021). Since cannibalistic morphs are more likely to injure kin due to possible physical proximity, inclusive fitness theory predicts that they should have more developed kin recognition abilities than non-cannibalistic morphs (Pfennig, 1999, 2021). Several theories such as theory of inclusive fitness (Penn and Frommen, 2010) and selfish gene (Gardner, and Welch, 2011) propose natural selection should favour individuals who can recognize their kin over those who cannot so that copies of individuals who can recognize their kin survive expanding the gene pool encoding this behaviour (Penn and Frommen, 2010; Clune *et al*., 2011; Mateo, 2015).

Kin recognition has been largely studied in eusocial insects with castes of soldiers or guards, *e*.*g*. ants, bees and termites (Lize *et al*., 2013; Vander Meer *et al*., 2019; Sinotte *et al*., 2021). Studies in desert isopods (*Hemilepistus reaumuri* Audouin and Savigny), paper wasps (*Ropalidia marginata* Lepeletier), and honeybees (*Apis mellifera* Linnaeus) have reported that they may use phenotypic cues or labels for discrimination between sibling and non-sibling conspecifics. More recently, Sohail *et al*. (2021) studied the cannibalistic expression of larval instar in green lacewing, *Chrysoperla carnea* Stephens (Neuroptera: Chrysopidae) towards related and unrelated conspecific eggs and reported that the larvae were more cannibalistic towards unrelated conspecific eggs and the rate of cannibalism increased in presence of conspecifics in the vicinity (Sohail *et al*., 2021). Also, adult *Drosophila melanogaster* Meigen is reported to have kin-recognition abilities based on specific cues associated with gut microbiome (Lewis *et al*., 2014; Lize *et al*., 2014; Carazo *et al*., 2015).

Coccinellids lay eggs in aggregative clusters in areas with high aphid density, and thus there is a risk of existence of overlapping stages in a given time and space, leading to increased competition between conspecifics over shared resources (Agarwala and Dixon, 1993a; Hodek *et al*., 2012). The egg laying females can be single or multiply mated and thus the cluster might consist of a mixture of sibling, half-sibling and non-sibling eggs. However, the time frame in which different stages coexists is short. It is highly likely that both larvae, as well as adults, will encounter sibling, half-sibling and non-sibling eggs. In addition, multiple females lay eggs in nearby location making it potentially difficult to identify between related and unrelated conspecifics. In this situation, there are chances that they utilise sensory information to avoid cannibalising their kin. Females of *Adalia bipunctata* Linnaeus (Agarwala and Dixon, 1993b) and *Propylea dissecta* Mulsant (Pervez and Khan, 2021) are able to recognise and avoid cannibalising their own eggs. In addition, larvae of *Harmonia axyridis* Pallas (Joseph *et al*., 1999), *A. bipunctata* (Agarwala and Dixon, 1993b), *P. dissecta* and *Coccinella transversalis* Fabricius (Pervez *et al*., 2005) are also reported to have kin recognition abilities through endogenous or chemical cues.

Based on literary background, cannibalism of eggs with varying degrees of relatedness (sibling, half-sibling, and non-sibling) by larval and adult stages was tested experimentally to determine whether *M. sexmaculatus* recognize siblings or not. It was hypothesised that the relatedness of larval and adult stages with the victim will modulate their cannibalistic tendency. Cannibals will recognize and avoid cannibalising sibling eggs in order to maximise their inclusive fitness.

## Materials and methods

### Stock culture

Adults of *Menochilus sexmaculatus* (*n*=60) were collected from the local agricultural fields of Lucknow, India (26°50’N, 80°54’E). The species was selected as an experimental model due to its abundance in local fields, wide prey range, and high reproductive output (Omkar et al., 2005). Adults were fed with *ad libitum* supply of cowpea aphid, *Aphis craccivora* Koch (Hemiptera: Aphididae). The aphid colonies were established on *Vigna unguiculata* L. plants in glasshouse (25 ± 2°C temperature, 65 ± 5% Relative Humidity). Adults were paired and placed in plastic Petri dishes (hereafter, 9.0 × 2.0 cm), which were kept in Biochemical Oxygen Demand incubators (Yorco Super Deluxe, YSI-440, New Delhi, India) at 25 ± 1°C, 65 ± 5% R.H., 14L: 10D. Eggs laid were collected daily, and held in plastic Petri dishes until hatching, which usually occurs within 2-3 days from oviposition. First instars were gently removed using a fine camel-hair paintbrush and assigned individually to clean experimental Petri dishes (size as above) once they began moving on or away from the remnants of their egg clutch.

### Collection of eggs used in choice treatment

For generation of sibling, half sibling and non-siblings, adults were randomly selected from stock culture and paired in different treatments as described below. For the production of siblings, reproductively mature, virgin and unrelated males and females were selected from the stock culture and paired in Petri dishes; one pair per dish. Post mating, males were removed and females were allowed to lay eggs. Eggs collected from females (family I) were divided into two groups, first group was used as experimental replicate and other was used as sibling eggs to be provided as victims in choice treatment.

For generation of half siblings, same males (family I) that mated earlier with the females (family I) were again mated with unmated, unrelated females (family II) collected from stock culture. The eggs obtained from these females (family II) were marked as half-sibling eggs and were further used in choice experiment.

For non-sibling eggs, unrelated males and females (family III) from different sub populations of stock culture were mated and eggs collected from these females were marked as non-sibling eggs. Fresh eggs were collected daily from respective females and were marked with their family code and relatedness for further use in the experiment as victims.

### Experimental setup

Different life stages were collected from family I, *i*.*e*. first, second, third, fourth instars and adult (*n*=15, each life stage) that were reared individually in Petri dishes on *ad libitum* supply of *A. craccivora*. At the start of the experiment, one experimental individual (any immature or adult stage) was placed in the experimental Petri dish containing three equidistantly placed clusters of twenty sibling, twenty half-sibling and twenty non-sibling eggs. The first victim cannibalised, time taken to encounter first victim, time taken for first consumption and first victim cannibalised were recorded for each life stage (first, second, third and fourth instar and adult). The total amount of eggs of differently related eggs, *i*.*e*. sibling, half-sibling, and non-sibling, cannibalised by each life stage was also recorded after 24 hours. For recording total amount of eggs cannibalised, the number of eggs provided were stage-specific, *i*.*e*. 20 eggs for first, 40 eggs for second, 60 eggs for third, 80 eggs for fourth and 100 eggs for adults of each sibling, half-sibling, and non-sibling eggs. The study was replicated fifteen times for each life stage, *i*.*e*. first, second, third and fourth instars and adults.

## Statistical analysis

To analyse the effect of victim relatedness on cannibalistic preferences of different life stages of *M. sexmaculatus*, victim first cannibalised by each life stage (larval instars and adults) were subjected to Chi-square test using Minitab 20.3 statistical software. Data sets on encounter time, latency to cannibalise and percent egg cannibalised were analysed with Shapiro-Wilk’s and Levene’s tests to test for normal distribution and variance homogeneity, respectively.

Further, to analyse the effect of victim encountered on time of first encounter, encounter time was used as response factor and life stage of the cannibal and victim encountered as fixed factors in a Generalised Linear Model (GLM).

To analyse the effect on latency to cannibalise, the latency to first victim cannibalised was used as response factor and life stage and victim cannibalised as well as their interaction as fixed factors in a GLM. For percent consumption, the data on percent victim cannibalised were used as response factor, and life stage and relatedness as well as their interaction as fixed factors in a GLM.

All the analyses were conducted using the Minitab 20.3 statistical software.

## Results

Chi-square analysis revealed insignificant effect of relatedness on the nature of victim first cannibalised by different life stages (χ^2^=1.92, P>0.05, df=8). In both larval stages as well as adults, victim first cannibalised was random (Figure 1).

**Figure 1.**
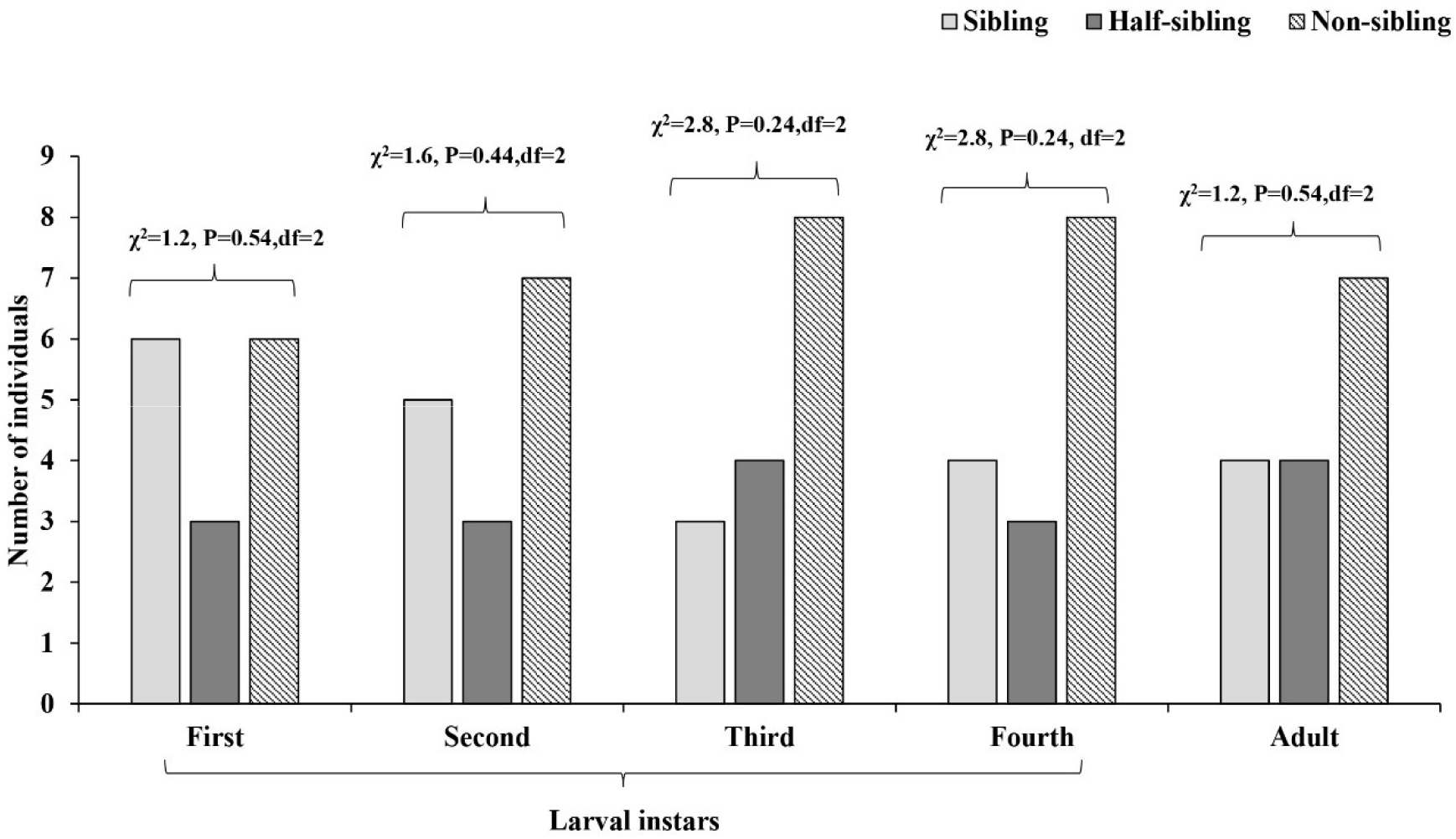
Effect of relatedness on first victim cannibalised by different larval instars and adults of *M. sexmaculatus*.

The time of first encounter was significantly different for different life stages (F=19.74, P<0.05, df=4,74) but was not affected by the relatedness of the victim cannibalised (F=0.36, P>0.05, df=2,74). The longest encounter time was recorded for first instars followed by third instars, fourth instars, adult and second instars (Figure 2).

**Figure 2.**
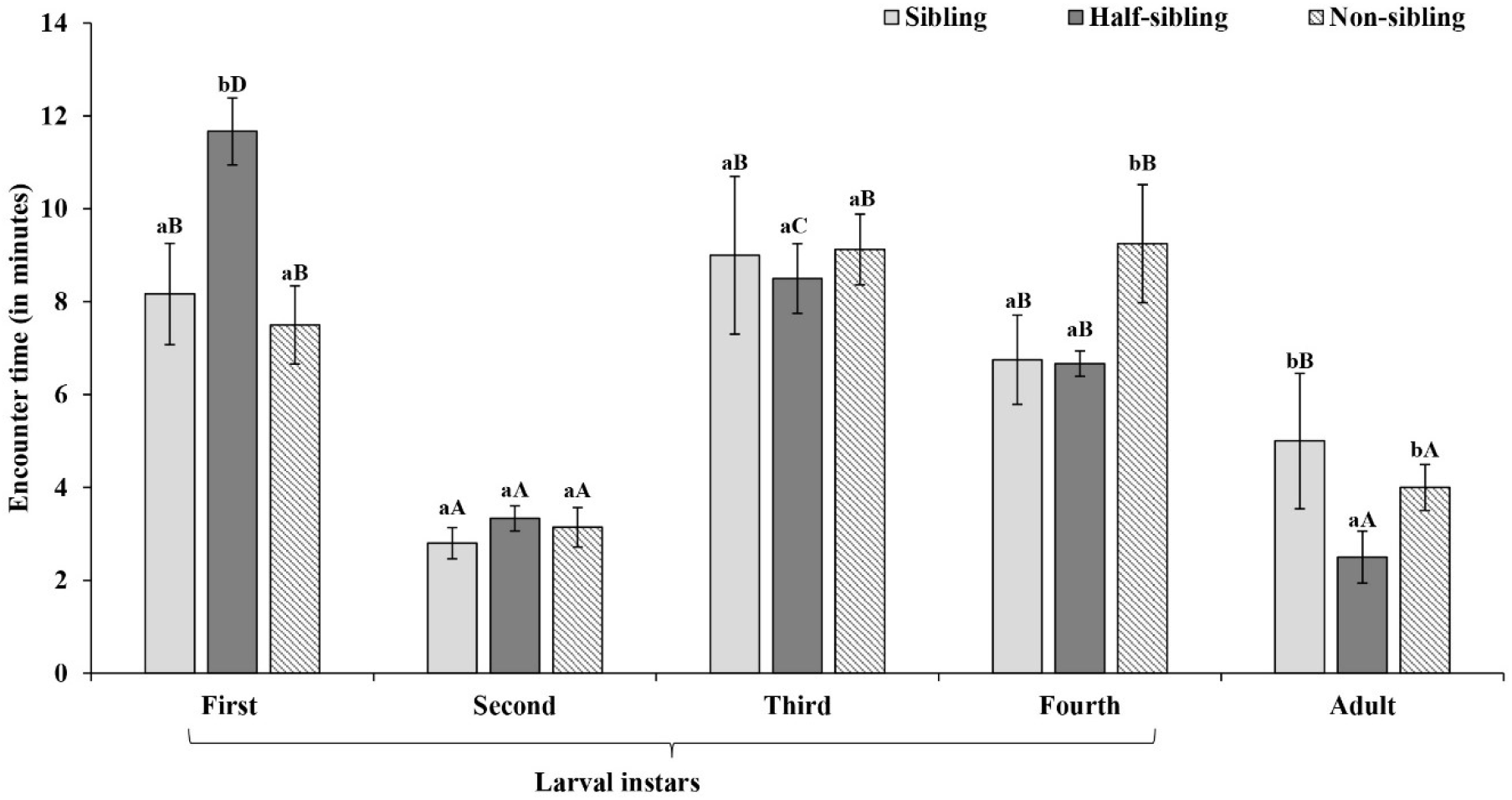
Effect of victim relatedness on encounter duration by different larval stages and adults of *M. sexmaculatus*. Values are mean ± SE. Lowercase and uppercase letters indicate comparison of mean within and between treatments respectively. Similar letters indicate lack of significant difference (P value > 0.05).

Similarly, the time of first victim cannibalised was significantly influenced by life stages (F=18.47, P<0.05, df=4,74). However, the consumption duration was not significantly affected by the relatedness of the victim cannibalised (F=1.83, P>0.05, df=2,74). Shortest consumption durations were recorded for first instars followed by third instars, fourth instars, second instars and adults (Figure 3).

**Figure 3.**
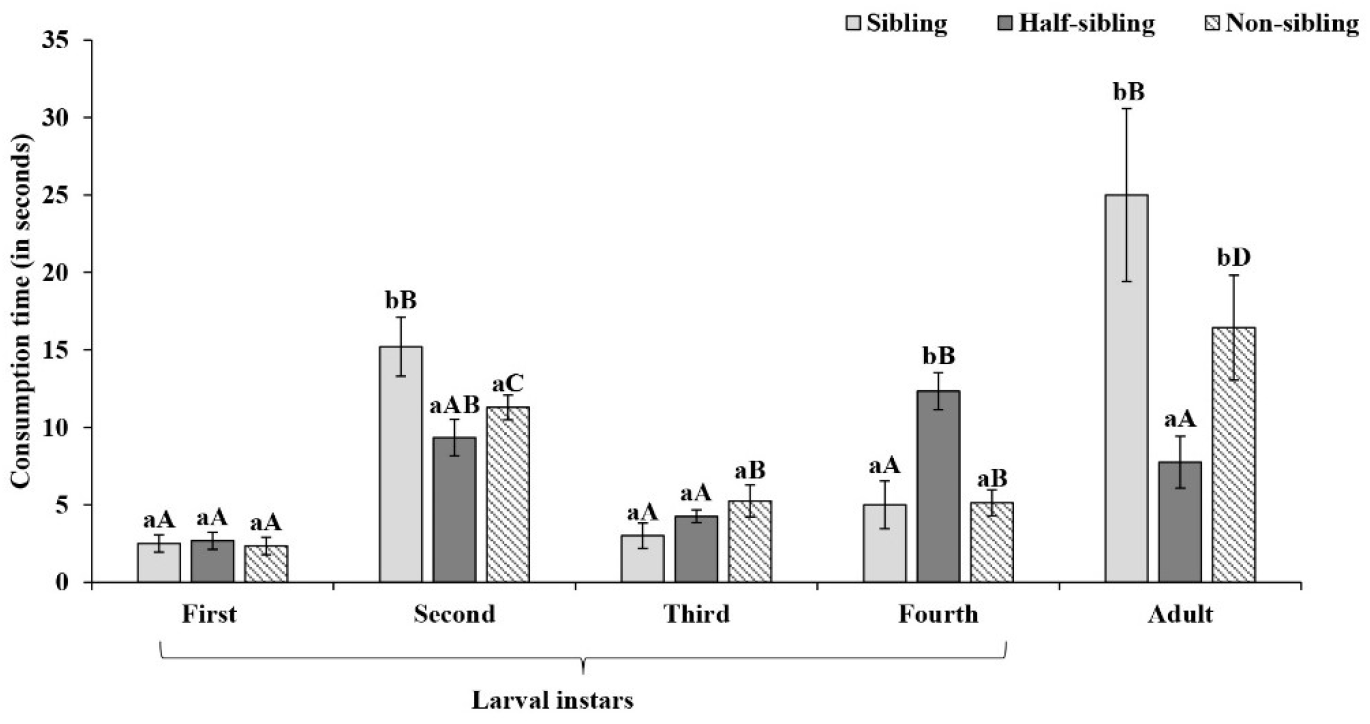
Effect of victim relatedness on consumption duration by different larval stages and adults of *M. sexmaculatus*. Values are mean ± SE. Lowercase and uppercase letters indicate comparison of mean within and between treatments respectively. Similar letters indicate lack of significant difference (P value > 0.05).

Percent eggs cannibalised by different life stages was significantly affected by both life stage (F=60.46, P<0.05, df=5,149) and relatedness (F=12.49, P<0.05, df=2,149) of the victim. In addition, their interactions were also found to be significant (F=2.74, P<0.05, df=8,149). Comparison of means on life stages revealed highest percent egg cannibalism by fourth instars followed by adults, third instars, second instars, and first instars.

In addition, comparison of means on relatedness revealed that the first instars tend to cannibalise more percentage of sibling and non-sibling eggs while second and third instars cannibalised more percentage of non-sibling eggs. Fourth instars and adults, on the other hand, cannibalised highest percentage of eggs irrespective of their relatedness (Figure 4).

**Figure 4.**
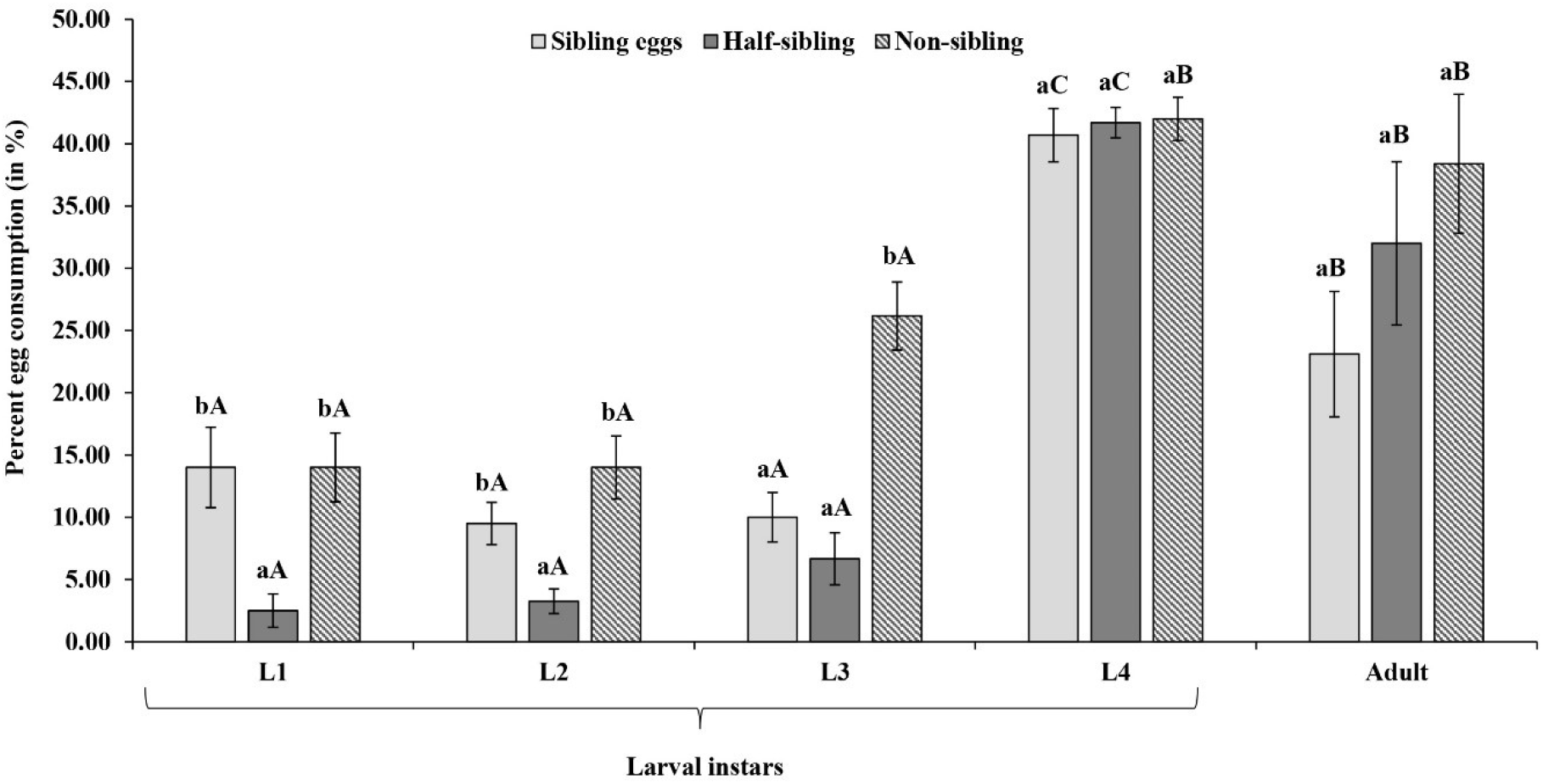
Effect of victim relatedness on percent egg consumption by different larval stages and adults of *M. sexmaculatus*. Values are mean ± SE. Lowercase and uppercase letters indicate comparison of mean within and between treatments respectively. Similar letters indicate lack of significant difference (P value > 0.05).

## Discussion

Current study revealed that the victim relatedness had insignificant effect on cannibalism by different larval instars and adults *M. sexmaculatus*. First encounter duration and the latency to cannibalise victim were found to be insignificantly affected by victim relatedness, however, both significantly varied with stage of the cannibal. Encounter duration decreased with the advancing stage except for the second instars and cannibalistic latencies followed a reverse trend. The victim first cannibalised by larval stages and adults were random, however, percent total egg consumption increased with advancing stage.

Insignificant difference was observed in victim first cannibalised by different larval instars and adults of *M. sexmaculatus*. However, significant differences in total percent number of eggs (after 24 hours) cannibalised with varying degree of victim relatedness suggests the presence of stage-specific cannibalistic tendencies and kin recognition mechanism in *M. sexmaculatus*. The percentage of total eggs cannibalised increased with the advancement in stage. First and second instars cannibalised a higher percentage of both sibling and non-sibling eggs while third instars cannibalised a higher percentage of non-sibling eggs. Fourth instars, and the adults, on the other hand, cannibalised the eggs regardless of their relatedness with the victim suggesting that kin recognition changes with stage. For first instars, mobility and the ability to tolerate hunger are relatively low, which might be a reason for high levels of sibling egg cannibalism (Ferran and Dixon, 1993). In addition, earlier studies have reported that the larval instars and adults can assess the surface chemical profile (Agarwala *et al*, 1998; Omkar *et al*., 2004). In *Hippodamia variegata* Goeze (Xie *et al*., 2022) and *H. axyridis* (Rondoni *et al*., 2021), antennal transcriptomes have reported the presence of odorant receptors and their chemosensory role in prey recognition and searching behaviour. Studies involving the role of family-specific chemical profiles in kin recognition have also been reported (Wong *et al*, 2014; Weiss and Schneider, 2021). Thus, the first instars cannibalising the eggs first encountered either sibling or non-sibling eggs might be attributed to the recognition of similar egg surface chemicals that lowers the risk of consuming toxic food and increases the chances of survival by overcoming the initial period of vulnerability, however, it does not confirm the presence of kin recognition ability in any of the life stages except the third instars which cannibalised higher percentage of non-sibling eggs. Previous investigations in *P. dissecta* and *C. transversalis* have also shown stage-specific cannibalistic responses towards sibling and non-sibling larvae, where third instars avoided sibling cannibalism while fourth instars indiscriminately cannibalised both sibling and non-sibling eggs (Pervez *et* al., 2005). Joseph *et al*. (1999) in a study on *H. axyridis* have reported that third instars avoided cannibalism of related victims and took longer and showed higher encounter rates to cannibalise related victims than unrelated victims. The highest percentage of eggs cannibalised by fourth instars may be attributed to increased evolutionary survival pressure and higher energetic needs required for pupation (Khan *et al*., 2003; Jafari, 2012; Khan and Yoldas, 2018a, b). The results suggest that possibly the nutritional requirements of larval instars vary based on the developmental stage which in turn regulates predatory prey consumption. In contrast, Agarwala and Dixon (1993b) in a study on *A. bipunctata*, have reported that female and young larvae are able to recognize kin, however, third instars showed no reluctance to eat second instar siblings.

Furthermore, the first encounters with the victim and the first cannibalistic attack on the victim by each life stage were independent of the degree of relatedness, indicating that these first encounters and consumptions were random events. However, both encounter duration and latency to cannibalise were significantly different for different life stages of the cannibal, suggesting that different life stages took different time durations before cannibalising eggs. Also, the differences in encounter durations among different life stages may be attributed to the extent of the mobility of the life stages and their need to procure food. Previous studies have shown that larval stages are known to alter their movement patterns following feeding as well as the area they transverse per unit time based on their size and age (Ferran and Dixon, 1993). For instance, the first instars would confine their search area to their immediate vicinity since they are the most critical developmental stage and with limited mobility, they typically stay close to the egg clutch. Post hatching, they first feed on their egg case and later on their neighbouring unhatched sibling eggs, which provide them with the energy necessary to search for food (Dixon, 2000; Hodek *et al*., 2012).

In conclusion, larval instars and adults prefer to cannibalise non-sibling eggs in *M. sexmaculatus*. The presence of kin recognition mechanism and discrimination among sibling and non-sibling eggs might play a beneficial role through increasing the inclusive fitness by decreasing the cannibalistic incidences among siblings and percent egg consumption increases with advancement of stage owing to the increased nutritional requirement and survival pressure.

## Acknowledgments

TY gratefully acknowledges CSIR, New Delhi, India, for Senior Research Fellowship, (09/107(0405)/2019-EMR-I) dated October 20, 2020. GM is thankful to Department of Higher Education, Government of Uttar Pradesh, India for providing financial assistance under the Centre of Excellence programme.

## Data availability statement

The datasets generated during and/or analysed during the current study are available from the corresponding author on reasonable request.

## References

Agarwala, B. K., & Dixon, A. F. G. (1993a). Kin recognition: egg and larval cannibalism in Adalia bipunctata (Coleoptera: Coccinellidae). European Journal of Entomology, 90, 45–50.

Agarwala, B. K., & Dixon, A. F. G. (1993b). Why do ladybirds lay eggs in clusters?. Functional Ecology, 541-548.

Agarwala, B. K., Bhattacharya, S., & Bardhanroy, P. (1998). Who eats whose eggs? intra- versus inter-specific interactions in starving ladybird beetles predaceous on aphids. Ethology Ecology and Evolution, 10(4), 361–368.

Anten, N. P. R., & Chen, B. J. W. (2021). Detect thy family: Mechanisms, ecology and agricultural aspects of kin recognition in plants. Plant Cell and Environment, 44(4), 1059–1071.

Bortolotti, G. R., Wiebe, K. L., & Iko, W. M. (1991). Cannibalism of nestling American kestrels by their parents and siblings. Canadian Journal of Zoology, 69(6), 1447–1453.

Bourke, A. F. G. (2014). Hamilton’s rule and the causes of social evolution. Philosophical Transactions of the Royal Society B: Biological Sciences. 369(1642), 20130362.

Carazo, P., Perry, J. C., Johnson, F., Pizzari, T., & Wigby, S. (2015). Related male Drosophila melanogaster reared together as larvae fight less and sire longer lived daughters. Ecology and Evolution, 5(14), 2787–2797.

Chiu, J. M. Y., Shin, P. K. S., Wong, K. P., & Cheung, S. G. (2010). Sibling cannibalism in juveniles of the marine gastropod Nassarius festivus (Powys, 1835). Malacologia, 52(1), 157–161.

Chung, M., Wang, M. Y., Huang, Z., & Okuyama, T. (2020). Diverse sensory cues for individual recognition. Development Growth and Differentiation, 62(9), 507–515.

Clune, J., Goldsby, H. J., Ofria, C., & Pennock, R. T. (2011). Selective pressures for accurate altruism targeting: Evidence from digital evolution for difficult-to-test aspects of inclusive fitness theory. Proceedings of the Royal Society B: Biological Sciences, 278(1706), 666–674.

Dixon, A. F. G. (2000). Insect predator–prey dynamics, ladybird beetles and biological control. Cambridge University Press, Cambridge, UK.

Dobler, R., & Kölliker, M. (2010). Kin-selected siblicide and cannibalism in the European earwig. Behavioral Ecology, 21(2), 257–263.

Dugas, M. B., McCormack, L., Gadau, A., & Martin, R. A. (2016). Choosy cannibals preferentially consume siblings with relatively low fitness prospects. American Naturalist.

Ferran, A., & Dixon, A. F. G. (1993). Foraging behaviour of ladybird larvae (Coleoptera: Coccinellidae). European Journal of Entomology, 90(4), 383–402.

Gardner, A., & Welch, J.J. (2011). A formal theory of the selfish gene. Journal of Evolutionary Biology, 24(8), 1801–13.

Gilbert, J. J. (2017). Non-genetic polymorphisms in rotifers: Environmental and endogenous controls, development, and features for predictable or unpredictable environments. Biological Reviews, 92(2), 964–992.

Henkel, S., & Setchell, J. M. (2018). Group and kin recognition via olfactory cues in chimpanzees (Pan troglodytes). Proceedings of the Royal Society B: Biological Sciences, 285(1889).

Hodek, I, Emden, H. F., & Honek, I. (2012). Ecology and behaviour of the ladybird beetles (Coccinellidae). Oxford, United Kingdom: Wiley-Blackwell.

Jafari, R. (2012). Cannibalism in Hippodamia variegata Goeze (Coleoptera: Coccinellidae) under laboratory conditions. International Symposium: Current Trends in Plant Protection Proceedings, 544–549.

Johnson, J. C., Kitchen, K., & Andrade, M. C. B. (2010). Family affects sibling cannibalism in the black widow spider, Latrodectus hesperus. Ethology, 116(8), 770–777.

Joseph, S. B., Snyder, W. E., & Moore, A. J. (1999). Cannibalizing Harmonia axyridis (Coleoptera: Coccinellidae) larvae use endogenous cues to avoid eating relatives. Journal of Evolutionary Biology, 12(4), 792–797.

Kalamara, M., Spacapan, M., Mandic-Mulec, I., & Stanley-Wall, N. R. (2018). Social behaviours by Bacillus subtilis: quorum sensing, kin discrimination and beyond. Molecular Microbiology, 110(6), 863–878.

Khan, M. H., & Yoldaş, Z. (2018). Investigations on the cannibalistic behavior of ladybird beetle Coccinella septempunctata L. (Coleoptera: Coccinellidae) under laboratory conditions. Turkish Journal of Zoology, 42(4), 432–438.

Khan, M. R., Khan, M. R., & Hussein, M. Y. (2003). Cannibalism and Interspecific Predation in Ladybird Beetle Coccinella septempunctata (L.) (Coleoptera: Coccinellidae) in Laboratory. Pakistan Journal of Biological Sciences, 6(24), 2013–2016.

Levis, N. A., & Ragsdale, E. J. (2022). Linking Molecular Mechanisms and Evolutionary Consequences of Resource Polyphenism. Frontiers in Integrative Neuroscience, 16.

Lewis, Z., Heys, C., Prescott, M., & Lizé, A. (2014). You are what you eat: Gut microbiota determines kin recognition in Drosophila. Gut Microbes, 5(4), 541–543.

Lightfoot, J. W., Dardiry, M., Kalirad, A., Giaimo, S., Eberhardt, G., Witte, H., & Sommer, R. J. (2021). Sex or cannibalism: Polyphenism and kin recognition control social action strategies in nematodes. Science Advances, 7(35).

Liu, X., Xia, J., Pang, H., & Yue, G. (2017). Who eats whom, when and why? Juvenile cannibalism in fish Asian seabass. Aquaculture and Fisheries, 2(1), 1–9.

Lizé, A., McKay, R., & Lewis, Z. (2013). Gut microbiota and kin recognition. Trends in Ecology and Evolution, 28(6), 325–326.

Lizé, A., McKay, R., & Lewis, Z. (2014). Kin recognition in Drosophila: The importance of ecology and gut microbiota. ISME Journal, 8(2), 469–477.

Mateo, J. M. (2015). Perspectives: Hamilton’s legacy: Mechanisms of kin recognition in humans. Ethology, 121(5), 419–427.

Mathiron, A. G. E., Pottier, P., & Goubault, M. (2019). Keep calm, we know each other: kin recognition affects aggressiveness and conflict resolution in a solitary parasitoid. Animal Behaviour, 151, 103–111.

Miranda, R. M., Espinoza, V., Dörner, J., Farías, A., & Uriarte, I. (2011). Sibling cannibalism on the small octopus Robsonella fontaniana (d’Orbigny, 1834) paralarvae. Marine Biology Research, 7(8), 746–756.

Modanu, M., Li, L. D. X., Said, H., Rathitharan, N., & Andrade, M. C. B. (2014). Sibling cannibalism in a web-building spider: Effects of density and shared environment. Behavioural Processes, 106, 12–16.

Omkar, & Bind, R. B. (2004). Prey quality dependent growth, development, and reproduction of a biocontrol agent, Cheilomenes sexmaculata (Fabricius) (Coleoptera: Coccinellidae). Biocontrol Science and Technology, 14, 665–673.

Omkar, Pervez A., Gupta, A. K., Pervez, O. A., & Gupta, A. K. (2004). Role of surface chemicals in egg cannibalism and intraguild predation by neonates of two aphidophagous ladybirds, Propylea dissecta and Coccinella transversalis. Journal of Applied Entomology, 128(9–10), 691–695.

Park, S., Jeong, J., & Park, D. (2005). Cannibalism in the Korean salamander (Hynobius leechii : Hynobiidae, caudata, amphibia) larvae. Integrative Biosciences, 9(1), 13–18.

Pener, M. P., & Simpson, S. J. (2009). Locust Phase Polyphenism: An Update. Advances in Insect Physiology, 36, 1–272.

Penn, D. J., & Frommen, J. G. (2010). Kin recognition: An overview of conceptual issues, mechanisms and evolutionary theory. Animal Behaviour: Evolution and Mechanisms, 55–85.

Pereira, L. S., Agostinho, A. A., & Winemiller, K. O. (2017). Revisiting cannibalism in fishes. Reviews in Fish Biology and Fisheries. Springer International Publishing.

Pervez, A., & Khan, M. (2021). Kin recognition by the adults of a biological control agent, Propylea dissecta (Coleoptera: Coccinellidae). Journal of Biological Control, 34(3), 227230.

Pervez, Ahmad, Gupta, A. K., & Omkar. (2005). Kin recognition and avoidance of kin cannibalism by the larvae of co-occurring ladybirds: A laboratory study. European Journal of Entomology, 102(3), 513–518.

Pfennig, D. W. (2021). Phenotypic Plasticity & Evolution Causes, Consequences, Controversies. Taylor & Francis.

Pfennig, D. W. (1999). Cannibalistic tadpoles that pose the greatest threat to kin are most likely to discriminate kin. Proceedings of the Royal Society B: Biological Sciences, 266(1414), 57–61.

Pfennig, D. W., & Collins, J. P. (1993). Kinship affects morphogenesis in cannibalistic salamanders. Nature, 362(6423), 836–838.

Pfennig, D. W., Reeve, H. K., & Sherman, P. W. (1993). Kin recognition and cannibalism in spadefoot toad tadpoles. Animal Behaviour, 46(1), 87–94.

Pfennig, David W., Sherman, P. W., & Collins, J. P. (1994). Kin recognition and cannibalism in polyphenic salamanders. Behavioral Ecology, 5(2), 225–232.

Rondoni, G., Roman, A., Meslin, C., Montagné, N., Conti, E., & Jacquin-Joly, E. (2021). Antennal transcriptome analysis and identification of candidate chemosensory genes of the harlequin ladybird beetle, Harmonia axyridis (Pallas) (coleoptera: Coccinellidae). Insects, 12(3), 1–16.

Rosati, G., Giari, A., & Ricci, N. (1988). Oxytricha bifaria (ciliata, hypotrichida): General morphology and ultrastructure of normal cells and giants. European Journal of Protistology, 23(4), 343–349.

Schutt, B. (2017). Eat me : a natural and unnatural history of cannibalism.

Singh, S., Mishra, G., & Omkar (2019). Oviposition in aphidophagous ladybirds: effect of prey availability and conspecific egg presence. International Journal of Tropical Insect Science, 39(2), 107–114.

Sinotte, V. M., Conlon, B. H., Seibel, E., Schwitalla, J. W., de Beer, Z. W., Poulsen, M., & Bos, N. (2021). Female-biased sex allocation and lack of inbreeding avoidance in Cubitermes termites. Ecology and Evolution, 11(10), 5598–5605.

Sohail, M., Soomro, Q. A., Asif, M. U., Rauf, I., & Muhammad, R. (2021). Conspecific neighbors and kinship influence egg cannibalism in the green lacewing, Chrysoperla carnea (Stephens). Egyptian Journal of Biological Pest Control, 31(1).

Soler, J. J., Martín-vivaldi, M., Nuhlícková, S., Ruiz-castellano, C., Mazorra-alonso, M., Martínez-renau, E., Svetlík, J., Eckenfellner, M., & Hoi, H. (2022). Avian sibling cannibalism : Hoopoe mothers regularly use their last hatched nestlings to feed older siblings. Zoological Research, 43(2), 265–274.

Tollrian, R. & Harvell, C. D., (1990). The evolution of inducible defence. (R. Tollrian & C. D. Harvell, Eds.), Princeton University Press (Vol. 100). Princeton University Press.

Uveges, B., Basson, A. C., Móricz, Á. M., Bókony, V., & Hettyey, A. (2021). Chemical defence effective against multiple enemies: Does the response to conspecifics alleviate the response to predators? Functional Ecology, 35(10), 2294–2304.

Vander Meer, R. K., Breed, M. D., Winston, M. L., & Espelie, K. E. (2019). Pheromone communication in social insects: ants, wasps, bees, and termites. CRC Press (Vol. 35).

Wall, D. (2016). Kin Recognition in Bacteria. Annual Review of Microbiology, 70, 143–160.

Walls, S. C., & Blaustein, A. R. (1995). Larval marbled salamanders, Ambystoma opacum, eat their kin. Animal Behaviour, 50(2), 537–545.

Weiss, K., & Schneider, J. M. (2021). Family-specific chemical profiles provide potential kin recognition cues in the sexually cannibalistic spider Argiope bruennichi. Biology Letters, 17(8), 20210260.

West, S. A., & Gardner, A. (2013). Adaptation and Inclusive Fitness. Current Biology, 23(13), R577–R584.

Wong, J. W. Y., Meunier, J., Lucas, C., & Kölliker, M. (2014). Paternal signature in kin recognition cues of a social insect: concealed in juveniles, revealed in adults. Proceedings of the Royal Society B: Biological Sciences, 281(1793), 20141236.

Xie, J., Liu, T., Yi, C., Liu, X., Tang, H., Sun, Y., & Zhang, Y. (2022). Antenna-Biased Odorant Receptor HvarOR25 in Hippodamia variegata Tuned to Allelochemicals from Hosts and Habitat Involved in Perceiving Preys. Journal of Agricultural and Food Chemistry. 70(4), 1090–1100.

Zaviezo, T., Soares, A. O., & Grez, A. A. (2019). Interspecific exploitative competition between Harmonia axyridis and other coccinellids is stronger than intraspecific competition. Biological Control, 131, 62–68.

